# Increased signal diversity/complexity of spontaneous EEG, but not evoked EEG responses, in ketamine-induced psychedelic state in humans

**DOI:** 10.1101/508697

**Authors:** Nadine Farnes, Bjørn E. Juel, André S. Nilsen, Luis G. Romundstad, Johan F. Storm

## Abstract

**Objective:** How and to what extent electrical brain activity is affected in pharmacologically altered states of consciousness, where it is mainly the phenomenological content rather than the level of consciousness that is altered, is not well understood. An example is the moderately psychedelic state caused by low doses of ketamine. Therefore, we investigated whether and how measures of evoked and spontaneous electroencephalographic (EEG) signal diversity are altered by sub-anaesthetic levels of ketamine compared to normal wakefulness, and how these measures relate to subjective assessments of consciousness.

**Methods:** High-density electroencephalography (EEG, 62 channels) was used to record spontaneous brain activity and responses evoked by transcranial magnetic stimulation (TMS) in 10 healthy volunteers before and after administration of sub-anaesthetic doses of ketamine in an open-label within-subject design. Evoked signal diversity was assessed using the perturbational complexity index (PCI), calculated from the global EEG responses to local TMS perturbations. Signal diversity of spontaneous EEG, with eyes open and eyes closed, was assessed by Lempel Ziv complexity (LZc), amplitude coalition entropy (ACE), and synchrony coalition entropy (SCE).

**Results:** Although no significant difference was found in the index of TMS-evoked complexity (PCI) between the sub-anaesthetic ketamine condition and normal wakefulness, all the three measures of spontaneous EEG signal diversity showed significantly increased values in the sub-anaesthetic ketamine condition. This increase in signal diversity also correlated with subjective assessment of altered states of consciousness. Moreover, spontaneous signal diversity was significantly higher when participants had eyes open compared to eyes closed, both during normal wakefulness and during influence of sub-anaesthetic ketamine doses.

**Conclusion:** The results suggest that PCI and spontaneous signal diversity may be complementary and potentially measure different aspects of consciousness. Thus, our results seem compatible with PCI being indicative of the brain’s ability to sustain consciousness, as indicated by previous research, while it is possible that spontaneous EEG signal diversity may be indicative of the complexity of conscious *content.* The observed sensitivity of the latter measures to visual input seems to support such an interpretation. Thus, sub-anaesthetic ketamine may increase the complexity of both the conscious content (experience) and the brain activity underlying it, while the *level, degree*, or general capacity of consciousness remains largely unaffected.

## 1 Introduction

“Consciousness” is often regarded as synonymous with “subjective phenomenological experience”, i.e. the experience of “what it is like” to have perceptions, feelings, and thoughts (Nagel, 1974). A challenging aspect of consciousness research is to define and assess different *levels* and *degrees* of consciousness. The concepts *level* and *content* of consciousness have been widely used and are often regarded as two orthogonal “dimensions” of consciousness, which may vary independently under some conditions (see Gress, 2009; Laureys, Boly, Moonen, & Maquet, 2009). Thus, the *level* of consciousness is commonly regarded as synonymous with degree of arousal or wakefulness, which is absent in coma, low during sleep, and high when we are wide-awake and alert (Laureys et al., 2009). In contrast, the *content* of consciousness – such as the richness of our experience - is thought to be variable within each such *level*. Thus, although conscious content and level normally appear to be correlated, there are exceptions such as the unresponsive wakefulness syndrome (UWS; also called “vegetative state”, VS) where there seems to be little or no discernible conscious content even during arousal (wakefulness). However, this scheme is not universally accepted. Some authors seem to use the term level differently (e.g. Casali et al, 2013), and some regard the level of arousal merely as a form of “background condition” that is necessary for consciousness to occur, rather than a dimension of consciousness itself (Koch, Massimini, Boly, & Tononi, 2016). Others still question the concept of levels of consciousness, and argue that the concept should be abandoned in exchange for a multidimensional account of global states of consciousness (Bayne, Hohwy, & Owen, 2016)

The integrated information theory of consciousness (IIT; Oizumi, Albantakis, & Tononi, 2014; Tononi & Edelman, 1998), developed by Giulio Tononi and colleagues, tries to characterize the necessary and sufficient properties required for any system to be conscious (Oizumi et al., 2014). IIT postulates that a system must be both integrated and differentiated to support consciousness (Oizumi et al., 2014). This prediction can to some extent be tested experimentally, albeit indirectly, in humans by use of the perturbational complexity index (PCI; Casali et al., 2013), which is assumed to assess the brain’s current capacity for sustaining consciousness by roughly indexing the brain’s potential to enter complex states, beyond ongoing activity. PCI is obtained by perturbing a small part of the cerebral cortex with transcranial magnetic stimulation (TMS) and measuring the resulting spatiotemporal pattern of electrical responses in large parts of the cortex with high-density electroencephalography (EEG). These responses are then analysed using the Lempel-Ziv compression algorithm to estimate the Kolmogorov complexity of the TMS-evoked activity (quantifying how compressible the signals are). Thus, PCI is a measure of the spatiotemporal complexity of global evoked cortical responses to a local perturbation, which is thought to reflect the complexity of the underlying system, including both its interconnectedness (integration), and the diversity of its available states of activity (differentiation).

So far, PCI has mainly been used to assess the brain’s capacity for complex dynamics in states that appear to differ drastically in terms of consciousness. For example, conscious states such as wakefulness and dreams, in which participants can confirm their subjective experience through some form of report, have been compared with apparently unconscious states such as dreamless sleep and some types of anaesthesia. PCI scores in healthy awake subjects are typically high, while PCI scores for participants in non-rapid eye movement (NREM) sleep and under general anaesthesia are low, irrespective of the location and intensity of the cortical TMS (Casali et al., 2013). The results indicate that the PCI scores during REM sleep (when conscious experience in the form of dreams often occur), in ketamine anaesthesia (which can cause vivid dreams), and in awake patients with locked-in syndrome (LIS, a condition where patients are fully conscious but unable to move or verbally communicate) are as high as for normal wakefulness (Casali et al.,2013). Thus, reduced PCI scores, indicating reduced integration and/or differentiation, seems to correlate well with reduced capacity for any form of experience (including dreaming and hallucinations), suggesting that PCI can be used as a general index of consciousness vs. unconsciousness (Casarotto et al. 2016).

Recent studies have indicated that neural differentiation alone, expressed as signal diversity in recordings of spontaneous brain activity, is consistently reduced in states typically associated with loss of consciousness (Schartner et al., 2015, Schartner, Pigorini, et al., 2017). In these studies, three measures of signal diversity were successfully applied to brain activity recordings; Lempel-Ziv complexity (LZc), amplitude coalition entropy (ACE), and synchrony coalition entropy (SCE). Compared to wakefulness, these signal diversity measures were found to be lower in states of apparent unconsciousness (sleep and propofol anaesthesia). More recently, the same researchers also found that signal diversity measured by magnetoencephalography (MEG) recordings of spontaneous brain activity, increased after giving doses of LSD, psilocybin and sub-anaesthetic ketamine compared to normal, resting wakefulness, suggesting that these psychedelic states are associated with an increase in neural differentiation (Schartner, Carhart-Harris, Barrett, Seth, and Muthukumaraswamy, 2017).

Like LSD and psilocybin, sub-anaesthetic doses of the dissociative anaesthetic ketamine, can cause psychedelic effects such as changes in perception and sensation, cognitive capacities, and experience of self, space, and time (Bowdle et al., 1998; Bayne & Carter, 2018). Sub-anaesthetic ketamine can therefore be used to investigate the neurobiological basis of a psychedelic state, a form of altered state of consciousness, while the subjects remain behaviourally responsive.

While the psychedelic state has been investigated with measures of neural differentiation, it seems important to measure PCI (which reflects both integration and differentiation) in psychedelic states of consciousness. This may shed light on the relationship between the capacity for consciousness, measured by combined integration and differentiation, and possible changes in the range of conscious content (Boly et al., 2013) that are thought to occur in psychedelic states (Gallimore, 2015). In contrast to signal diversity, which is considered to measure differentiation but not integration, PCI has to our knowledge not previously been tested in any psychedelic state.

Therefore, our main aim in this study was to investigate whether and how PCI is affected in a psychedelic state, induced by sub-anaesthetic ketamine, compared to normal wakefulness. In addition, we wanted to test whether the reported increase in signal diversity in spontaneous MEG signals during the psychedelic state (Schartner, Carhart-Harris, et al., 2017) would extend also to spontaneous EEG signals. Finally, we aimed to investigate the relationship between the subjectively reported effects of sub-anaesthetic ketamine and the signal diversity measures.

Results from this work were previously presented at two conferences in 2017 (Farnes et al. 2017a, Farnes et al. 2017b) and as a preprint manuscript in January 2019 (Farnes et al 2019).

## 2 Methods

This study was approved by the regional committees for medical and health research ethics (2015/1520/REK sør-øst A) and written informed consent was obtained from all participants before the start of the study. Participants were recruited by posters at the university campus and participants received financial compensation for participation in the study.

### 2.1 Participants

We recruited 34 participants. Participants were excluded if: (1) not healthy, (2) under the age of 18, (3) incompatibility with MRI scanning (metal or electric implants, pregnancy, breastfeeding, reduced kidney function and claustrophobia), (4) incompatibility with TMS administration (recent loss of consciousness caused by head injury or epilepsy), (5) incompatibility with ketamine administration (somatic diseases, previous or present neurological or psychiatric illnesses, psychiatric illnesses in family members, medication or allergies that could interact with ketamine, substance abuse, recent or regular drug use, previous adverse reaction to drugs, or needle phobia), (6) difficulty finding suitable resting motor threshold for TMS (RMT, i.e. the stimulation intensity at which 50% of TMS pulses over the optimal spot in primary motor cortex generate twitches in the pollicis brevis (thumb) muscle as recorded with surface electromyography (Rossini et al., 2015)), and, (7) low TMS evoked potential (TEP) quality (muscle artefacts and peak to peak amplitude less than 10 µV). 10 participants were included and completed the study.

### 2.2 Procedure

All experimental procedures were carried out at the Intervention Centre at Oslo University Hospital. Before the main experiment, spontaneous EEG and TMS evoked EEG responses were recorded without ketamine (day 1) to assess participants’ TEPs to ensure that only participants with strong responses to TMS completed the main experiment (day 2), about 6 days after day 1 (median 6 days; range: 3-23 days). Two TMS-EEG and spontaneous EEG sessions were completed on day 2; baseline and intervention involving administration of sub-anaesthetic doses of ketamine (see **Figure S1** for a flow chart of the stages in the study). On day 1, we first found the RMT and searched for an optimal target TMS stimulation location for the main experiment. Thereafter, 300 TMS-EEG trials were recorded while the participants were in a restful, wakeful state with their eyes open. In addition, two 2-minute segments of spontaneous EEG were recorded - first with eyes open and then with eyes closed. Thereafter, participants were given intravenous infusion of sub-anaesthetic doses of ketamine. When the participant reached a state in which they reported a noticeable effect of the drug, we measured the RMT again (Di Lazzaro et al., 2003), and adjusted the TMS stimulation intensity to the same percentage of the RMT as was used during the wakeful state. We then delivered another 300 TMS pulses, while keeping the stimulation target the same as before ketamine administration. Finally, spontaneous EEG was recorded with the same conditions as before ketamine administration.

During the stimulation period the participants wore earphones with a noise masking sound, made from randomly scrambled TMS clicks, to reduce auditory potentials evoked by the TMS clicks (Ilmoniemi & Kicic, 2010). The noise masking sound was adjusted so that the TMS click could not be heard but was never so loud that the sound became uncomfortable for the participant. Throughout the procedure, participants were lying down and had their head fixed on a vacuum pillow to ensure stability during stimulation.

#### 2.2.1 Identifying target area for TMS stimulation

On day 1, based on each participant’s MR image, a suitable target point for stimulation was identified within the parietal (Brodmann Areas: BA 7, BA 5) or prefrontal cortex (BA 6), similar to the procedures used in Casarotto et al. (2016) and Casali et al. (2013). If there were no large muscular or magnetic artefact lasting over 10 ms visible in the EEG response to single pulses, 20-30 pulses were given, and the resulting TEP was examined online. If the TEP amplitude was below 10 µV peak-to-peak within the first 50ms, we tried to improve the TEP signal by increasing the intensity. If this did not improve the TEP or introduced more artefacts, we adjusted coil rotation or position. The target point for stimulation was accepted if a non-artefactual TEP, with a peak-to-peak amplitude equal to or exceeding 10 µV in the channels near the stimulation site in the first 50 ms after stimulation, was observed in the averaged TEP signal following 30 consecutive single pulses. When the stimulation area was found, 300 single pulses were given with a random jittering inter-stimulus interval (range: 1.7 – 2.3 seconds; Casali et al. 2013). One participant was stimulated in BA 4 because this was the only area without large muscle artefacts (See Table S1 for an overview of the TMS targets and stimulation parameters for all participants). Navigation (Neuro-navigation with PowerMag View) of the coil position and angle relative to the participant’s brain was used to minimize the deviation from the set target point position. For increased efficiency the centre of the coil was positioned tangentially to the scalp (Rossi, Hallett, Rossini, Pascual-Leone, & Safety of T. M. S. Consensus Group, 2009). The procedure for finding an area of stimulation was the same for day 1 and day 2.

#### 2.2.2 Ketamine administration

Participants were told to abstain from food 6 hours before the first recording session started and from drinking 2 hours before ketamine administration. Racemic ketamine 10 mg/ml (Ketalar®, Pfizer AS, Lysaker, Norway) was administered by an anaesthesiologist or a nurse anaesthetist by continuous intravenous infusion using an infusion pump (Braun®) in increasing steps from 0.1 to 1.0 mg/kg/h, increasing with 0.1 mg/kg/h every fifth minute. The participants were asked to report when they felt they had a possible drug effect and when they were certain that they had an effect. When both the anaesthesiologist and the participant were certain that the participant had an effect of the drug, the continuous ketamine infusion rate was stabilized to maintain the subjective effect (B. Braun Perfusor Space, B. Braun Melsungen AG, Melsungen, Germany). The median continuous infusion was 0.7 mg/kg/hr (range: 0.5 to 1.0 mg/kg/hr) for each participant, producing psychotomimetic effects without loss of consciousness (Domino, Chodoff, & Corssen, 1965) See supplementary material for maintained, continuous dose, total dose received for each participant, and dosage steps over time. Participants’ pulse oximetry and heart rate was continuously monitored during the ketamine administration by the anaesthesiologist or the nurse anaesthetist. Median administration time was 43.5 minutes (range: 37 – 73 minutes). Participants could leave the hospital facilities after the anaesthesiologist had checked to ensure that the effects of the drug had subsided (approximately 2 hours after discontinuation of ketamine administration). A follow-up email was sent to the participants more than a week after the finished experiment to check their wellbeing.

#### 2.2.3 Psychedelic assessment: 11D-ASC

To assess the psychedelic effects of ketamine relative to the pre-ketamine condition, participants retrospectively rated the content of their experience using an altered states of consciousness questionnaire: an extended, 11-dimensional version (11D-ASC) of the original 5-Dimensional Altered States of Consciousness Rating Scale (5D-ASC; Dittrich, 1998, translated to English from German by Felix Hasler and Rael Cahn), translated into Norwegian. The 11D-ASC questionnaire has 11 subscales: 1. *experience of unity*, 2. *spiritual experience*, 3. *blissful state*, 4. *insightfulness*, 5. *disembodiment*, 6. *impaired cognition and control*, 7. *anxiety*, 8. *complex imagery*, 9. *elementary imagery*, 10. *audio-visual synaesthesia*, and 11. *changed meaning of percepts* (Studerus, Gamma, & Vollenweider, 2010). For each statement in the questionnaire, participants were told to indicate their level of agreement on a visual analog scale (VAS) anchored from “No, not more than usual” (left) to “Yes, much more than usual” (right), and scored by using percentage (left to right). An example of a statement is “I saw things I knew were not real”. Participants were instructed to respond considering the time interval from when they felt they had an effect of the drug until the effects subsided, and where the “usual state” was before ketamine administration. Thus, the score indicated strength of experience relative to the normal non-psychedelic state. The mean score of all the questions gives the Global-ASC score. The 11D-ASC was administered 45-60 minutes after discontinuation of ketamine after most drug effects had subsided.

### 2.3 Setup

Individual T1 weighted structural MR images (Phillips 3.0T Ingenia MR system, Philips Healthcare, The Netherlands) were obtained from each participant for spatial navigation to precisely locate the cortical target for TMS stimulation. For neuro-navigation we used the PowerMag View! system (MAG & More GmbH, München, Germany). This system uses two infrared cameras (Polaris Spectra) to track the position of the participant’s head and TMS coil in space. A figure-eight coil (Double coil PMD70-pCool, MAG & More GmbH, München, Germany) was used for stimulation (maximum field strength of 2T (∼ 210 V/m), pulse length of 100 µs, winding diameter of 70mm, biphasic pulse form) driven by a PowerMag Research 100 stimulator (MAG & More GmbH, München, Germany). The RMT was determined using PowerMAG Control (MAG & More GmbH, München, Germany). EEG was recorded with two 32-channel TMS-compatible amplifiers (BrainAmp DC, Brain Products, Germany) connected to a 60 channel TMS-compatible EEG cap. In addition, two electrodes detected eye movements (EOG), and a common reference was positioned at the forehead with the ground electrode. The impedance of all EEG electrodes was kept under 10 kΩ. EEG signals were sampled at 5000 Hz with 16-bit resolution and a 1000 Hz low pass filter was applied upon acquisition.

### 2.4 Analysis

#### 2.4.1 TMS-EEG pre-processing and PCI analysis

All pre-processing of the TMS-EEG data was done manually using the MATLAB (MATLAB R2016A, The Mathworks) based SiSyPhus Project (SSP 2.3e, University of Milan, Italy). EEG responses to TMS were visually inspected to identify artefactual trials and channels containing abnormal amplitude activity which were excluded from further analysis. The interval around the time of the TMS-pulse (−2ms to 5ms) was removed for all participants to exclude the TMS artefact, and artefactual channels (flat, noisy or with abnormally high amplitude over a large duration of the recording) were interpolated. Trials with abnormal voltage traces (high variance, large transient deflections, movement artefacts etc.) in multiple channels were rejected and removed from further analysis. The remaining data was zero-centered (mean baseline correction) to eliminate any voltage offsets. Any residual artefacts in the data after 5 ms was detected by independent component analysis (ICA) and then removed. Rejection of artefactual components was done manually by inspecting EEG component topography, activity over time and power spectrum. Eye blink, eye movement and other muscle and non-physiological artefacts as described in Rogasch et al. (2017) and Hamidi, Slagter, Tononi, and Postle (2010) were removed, while components that seemed to contain at least some physiological characteristics were kept (Rotenberg, Horvath, & Pascual-Leone, 2014). Signals were referenced to the common average reference, using a 1 Hz high pass filter and 45 Hz low pass filter (Hamming window sinc FIR filter), and downsampled to 312.5 Hz.

The remaining analyses used for PCI calculation were fully automatic and performed by use of MATLAB scripts (MATLAB R2013A, The Mathworks) courtesy of Adenauer Casali (University of Milan, Italy) as described in Casali et al., (2013). Source estimation of significant cortical sources was done by using a standard head model from the Montreal Neurological Institute (MNI) atlas. First, the significantly active cortical sources were estimated using a threshold set by the 99th percentile of the distribution of maximum amplitudes of bootstrap resampled baseline activities before TMS. If the amplitude of the source at a specific time exceeded this threshold, it was given the value 1, otherwise it was given the value 0. This resulted in a binarized matrix of significant sources over time, time-locked to the TMS pulse. Data in the interval 8-300 ms after the pulse of the resulting binarized matrix was used to derive the PCI value by calculating the Lempel-Ziv complexity (LZc), using the LZ76 compression algorithm. Finally, the LZc was normalized by the asymptotic maximum complexity of a matrix of the same size containing the same proportion of 1s to 0s, yielding the PCI value for the session (see supplementary material of Casali et al., 2013 for a detailed explanation).

To avoid instability in the PCI values, a threshold for source entropy was set to 0.08, in accordance with Casali et al. (2013). All of the TMS-EEG recordings exceeded this threshold (*mean* = 0.6, *SD* = 0.2) and all the data showed a high signal-to-noise ratio (>2, indicative of the EEG responses being closely related to the TMS stimulus, **Figure S2**).

#### 2.4.2 Spontaneous EEG preprocessing and signal diversity analysis

The spontaneous EEG data was pre-processed using EEGlab. First, the data was split into 8-second non-overlapping segments, resulting in 15 epochs per condition (normal wakefulness and during ketamine infusion, with eyes open and eyes closed). Then, artefactual channels were marked, removed, and interpolated. Epochs were rejected based on the same criteria as for the TMS-EEG data and the data was baseline corrected (zero mean). Signals were referenced to the common average reference, filtered with a 0.5 Hz high pass and 45 Hz low pass filter (Hamming window sinc FIR filter), and downsampled to 250 Hz. ICA components such as eye blinks and eye movement were manually detected and excised from the data. As in Schartner et al. (2015), the surface Laplacian of the data was computed, increasing topographical specificity by subtracting the averaged signal of each channels’ nearest neighbours (Hjorth, 1975). From the 62 channels, only 9 were chosen for signal diversity analysis (**Figure S2**) due to the entropy measures being calculated based on the distribution of states observed in the data. Since number of states, S, available in a network of N binary nodes is *S = 2*^*N*^, and the measures require an estimate of the probability density distribution over states, the number of samples in an epoch should be at least as large as the number of states available to yield a representative estimate of the underlying distribution of states. With each epoch containing 2000 samples (8s epoch length x 250Hz sampling rate), the maximal number of channels that could be included in the analysis if all states were to have a chance of being sampled at least once, would be 10. However, to increase the chance of getting a decent sampling of the distribution, we decided to use only 9 channels.

The signal diversity measures amplitude coalition entropy (ACE), synchrony coalition entropy (SCE), and Lempel Ziv Complexity (LZc), were calculated as described in Schartner et al. (2015). We first performed a Hilbert transformation and then binarized the data. The binarization threshold was set to the mean absolute amplitude (ACE, LZc) or to the absolute phase synchrony between each channel pair according to a 0.8 radian threshold (SCE). Next, we found the distribution of states over time, defining a state as a binary string of the activity over channels (ACE) or phase synchrony over channel pairs (SCE) at a given time point. Shannon entropy was then calculated over the state distributions and normalized according to the maximum entropy of a randomized sequence with similar characteristics as the original (ACE, SCE). Finally, the mean values were calculated over epochs (ACE) or channel pairs and epochs (SCE). LZc was calculated by directly applying the LZ76 algorithm (Kaspar & Schuster, 1987; Lempel & Ziv, 1976) to the spatially concatenated binarized activity matrix, calculating the compressibility of the data. Normalization was done by dividing the resulting raw value with the LZc of the same data shuffled in time.

#### 2.4.3 Statistical analysis

All statistical analyses were done in SPSS (IBM SPSS Statistics 24). To investigate significant differences between the sub-anaesthetic ketamine condition and the normal wakeful state on RMT and PCI we used the parametric paired-samples T-test for PCI, and the non-parametric Wilcoxon signed-rank (WSR) test for RMT. Normality was determined by the Shapiro-Wilk test (significance p > 0.05). To assess the effects of spontaneous signal diversity in sub-anaesthetic ketamine compared to normal wakefulness and eyes open compared to eyes closed, we used a repeated two-way ANOVA. A linear mixed model was used to assess whether changes in stimulation intensity affected the spatiotemporal activation values (average of the significant cortical sources activated after TMS) and PCI values. A random intercept was included to account for within-subject correlations. The model parameters were estimated using restricted maximum likelihood, and statistical significance was assessed by t-tests.

To assess significant relationships, the non-parametric Spearman’s rank order correlation was used. Correlations were assessed for the relationship between PCI values and stimulation intensity, PCI and spontaneous signal diversity measures and continuous and total ketamine dose, and total ketamine dose and global-ASC score. Moreover, PCI and spontaneous signal diversity in the ketamine condition was correlated with the 11-ASC and global-ASC. Significance threshold for all tests was p < 0.05.

## 3. Results

### 2.5 Increased spontaneous, but not evoked signal diversity in sub-anaesthetic ketamine compared to normal wakefulness

Our main aim was to investigate whether evoked signal diversity measured by PCI, but also spontaneous signal diversity measured by LZc, ACE, and SCE, is affected by sub-anaesthetic doses of ketamine when comparing to the normal wakefulness. For PCI, we first investigated the spatiotemporal response to TMS (**Figure 1**) and found that both in the normal wakeful state and the sub-anaesthetic condition, the brain responded to TMS with long-lasting patterns of activation. The response was not limited to the site of activation, as seen by the distribution of significant sources, but spread to different cortical locations. Qualitatively, the spatiotemporal characteristics appeared similar in both conditions, with small variations in amplitude and latencies between the two conditions for individual participants. In accordance with these observations, we found no significant difference between PCI values for normal wakefulness (*mean* = 0.53, *SE* = 0.02) and sub-anaesthetic ketamine (*mean* = 0.55, *SE* = 0.03, *t* (9) = −0.87, *p* = 0.41, *r* = 0.27, **Figure 2**).

**Figure 1.**
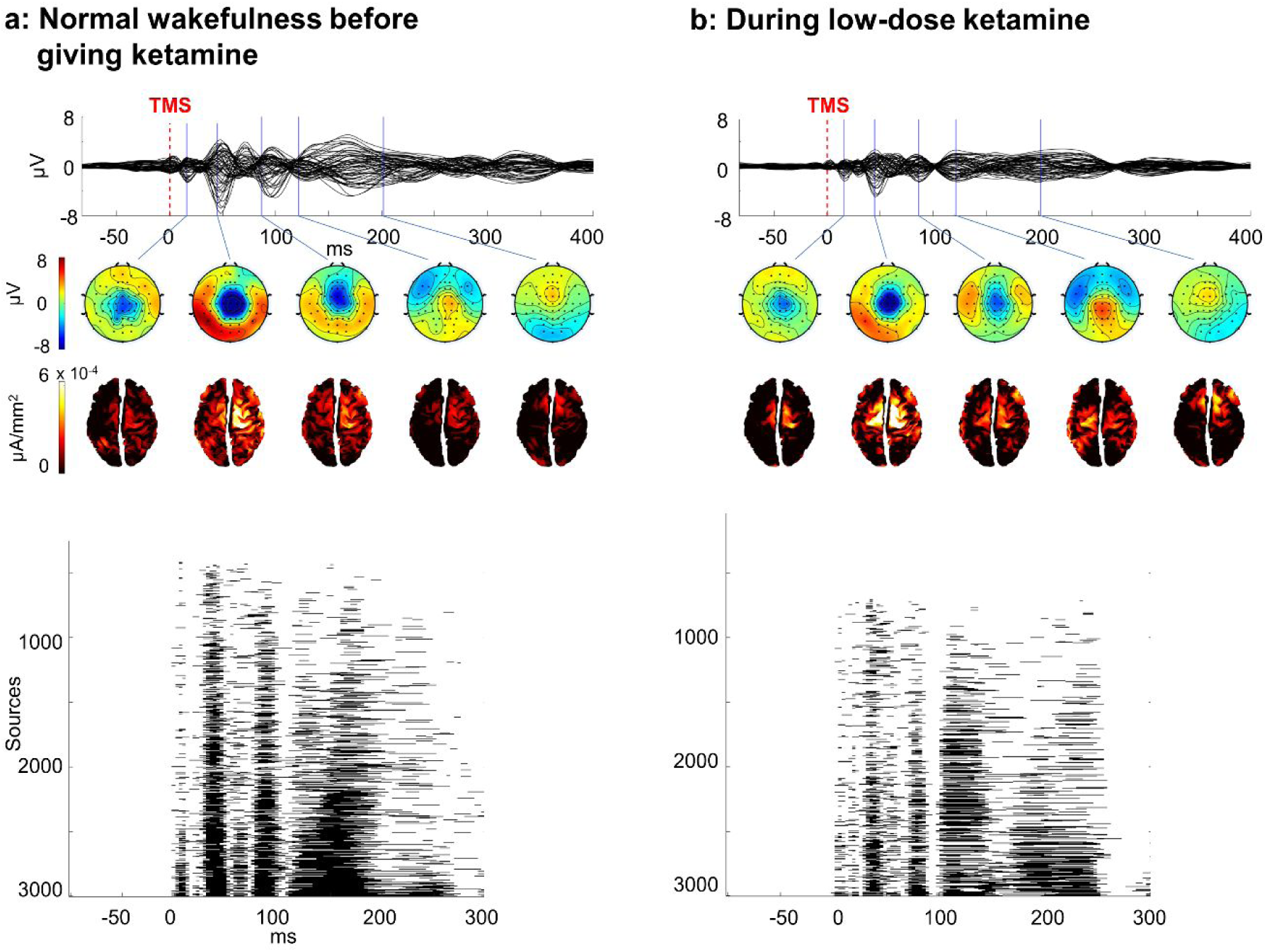
Spatiotemporal dynamics of TMS-EEG responses in normal wakefulness versus ketamine-induced psychedelic state. Averaged TMS-evoked potentials (298 and 281 trials) over all EEG channels in one representative subject before (**a**) and after (**b**) ketamine administration. The stimulation intensities were 80% and 79% of the maximal stimulator output, respectively. Underneath voltage topographies, reflecting the electrical activity across the scalp, and corresponding distributions of significant cortical currents are displayed at selected latencies. Underneath each figure are the binary SS(x,t)-matrices where significant sources at a given time are displayed as black, and white if not significant. The sources are ordered after total amount of significant activation in the response after TMS from bottom to top.

**Figure 2.**
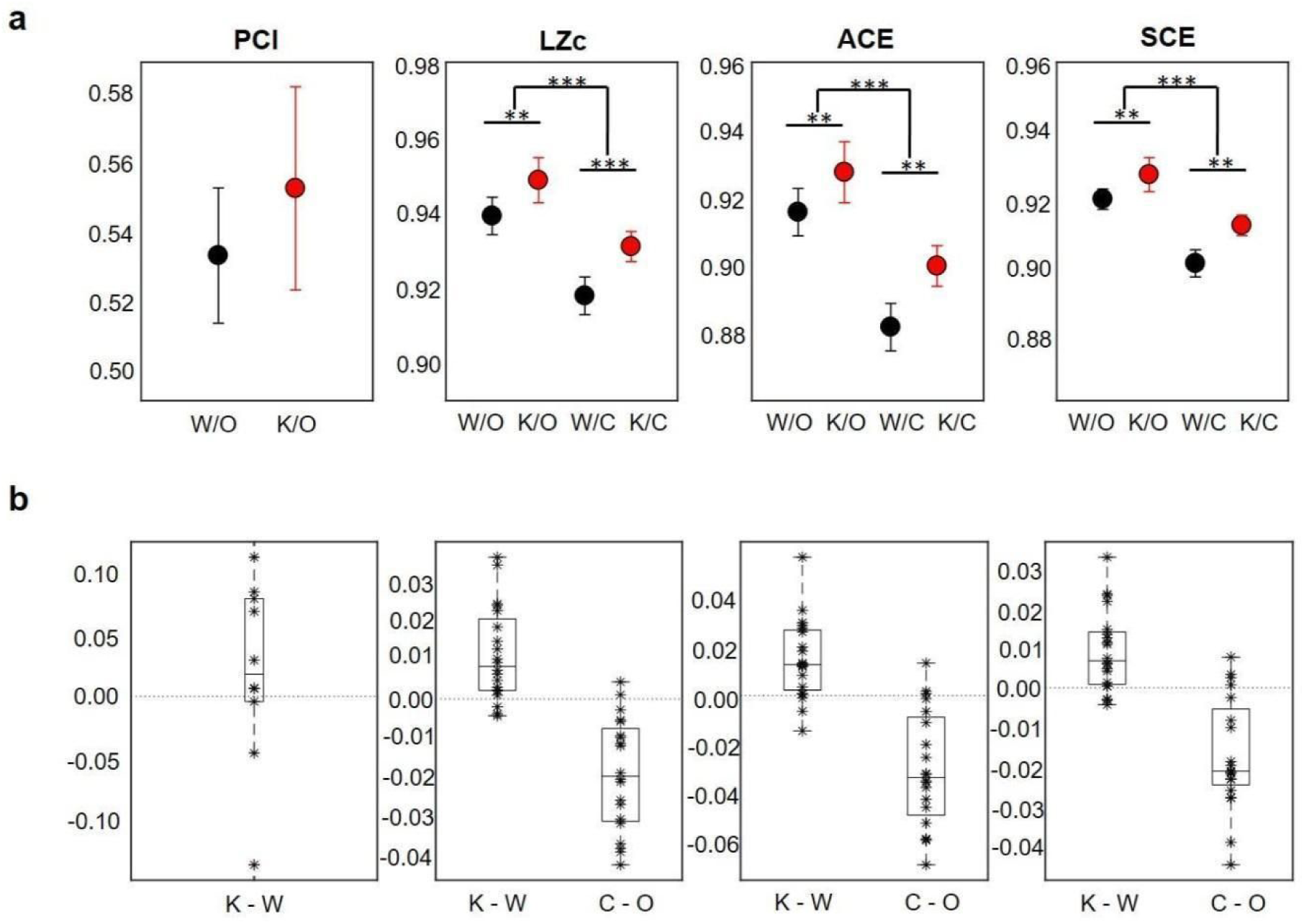
Average values and difference values for PCI, LZc, ACE, and SCE. **a**) Average values with one standard error of the mean (SEM) error bars for PCI, LZc, ACE, and SCE in wakefulness with eyes open (W/O) and eyes closed (W/C), and ketamine eyes open (K/O) and eyes closed (K/C). The stars (*, **, and ***) indicate statistical significance (p <0.05, p <0.01, p < 0.001) between wakefulness and ketamine, and eyes open or closed. **b**) Boxplots showing differences in individual PCI, LZc, ACE, and SCE values subtracting ketamine from wakefulness (K-W) and eyes closed from eyes open (C-O).

For all the spontaneous signal diversity measures (LZ, ACE, and SCE), we found a significant increase in the sub-anaesthetic ketamine condition compared to normal wakefulness (LZ: *F*(1,9) = 11.13, *p* < 0.05, *r* = 0.75, ACE: *F*(1,9) = 10.67, *p* < 0.05, *r* = 0.74 and SCE: *F*(1,9) = 11.79, *p* < 0.05, *r* = 0.75). Moreover, we found that all measures were significantly higher when participants had their eyes open compared to when they had their eyes closed (LZ: *F*(1,9) = 20.83, *p* < 0.05, *r* = 0.84, ACE: *F*(1,9) = 17.78, *p* < 0.05, *r* = 0.81, and SCE: *F*(1,9) = 16.71, *p* < 0.05, *r* = 0.81). No interaction effects between signal diversity before and after ketamine and eyes open/eyes closed for the signal diversity measures was observed.

### 2.6 Signal diversity values were unaffected by stimulation intensity and sub-anaesthetic ketamine dose

Since RMT was measured before and after sub-anaesthetic ketamine to determine the intensity for TMS stimulation, we wanted to investigate whether stimulation intensity affected the PCI values. First, we found no significant difference in RMT before (*median*: 54.5, *range*: 46.5 - 64) and after ketamine administration (*median*: 56.8, *range*: 43 - 62, *z* = −0.48, *p* = 0.64, *r* = −0.11). Moreover, the mean change in stimulation intensity relative to maximum stimulator output from before to after sub-anaesthetic ketamine was a 0.81 % decrease (*SD*: −9.7% – 14.6%), but this change in stimulation intensity did not significantly affect spatiotemporal activation values (*regression coefficient* [95 % CI] = −3×10^-3^, [-4×10^-3^ 12×10^-2^], *p* = 0.35), nor PCI values (*regression coefficient* [95 % CI] = −0.04, [-8×10^-3^ 0], *p* = 0.07). Intra-class correlation coefficients were found to be 0.88 for spatiotemporal activation and 0.31 for PCI.

For ketamine doses, we found no significant correlation between the rate of continuous maintained ketamine dose and PCI (*r*_*s*_ = −0.04, *p* = 0.90) or spontaneous signal diversity (LZc eyes open: *r*_*s*_ = 0.04, *p* = 0.92, LZc eyes closed: *r*_*s*_ = 0.05, *p* = 0.89, ACE eyes open: *r*_*s*_ = 0.05, *p* = 0.89, ACE eyes closed: *r*_*s*_ = −0.04, *p* = 0.92, SCE eyes open: *r*_*s*_ = 0.03, *p* = 0.93, SCE eyes closed: *r*_*s*_ = 0.20, *p* = 0.58). Similarly, we found no significant correlation between total ketamine doses and PCI values (*r*_*s*_ = 0.18, *p* = 0.63) or spontaneous signal diversity values (LZc eyes open: *r*_*s*_ = −0.03, *p* = 0.93, LZc eyes closed: *r*_*s*_ = 0.15, *p* = 0.68, ACE eyes open: *r*_*s*_ = 0.02, *p* = 0.96, ACE eyes closed: *r*_*s*_ = 0.10, *p* = 0.78, SCE eyes open: *r*_*s*_ = 0.07, *p* = 0.86, SCE eyes closed: *r*_*s*_ = 0.26, *p* = 0.47).

### 2.7 Correlations with alterations in phenomenology

All participants retrospectively reported to have had an effect of ketamine, but to differing degrees (**Figure 3A**). However, the overall average response to the 11D-ASC showed that the subscales disembodiment, complex imagery, and elementary imagery had the highest scores, while the subscale anxiety had the lowest score (**Figure 3B**). No statistical significant correlations were found between total ketamine dose and global-ASC scores (*r*_*s*_ = 0.33, *p* = 0.35). However, the subscales impaired cognition and control, anxiety and changed meaning of perception had high correlations (*r* > 0.5) with the signal diversity measures in the eyes open condition, while anxiety had the highest correlation in the eyes closed condition (**Figure 3C**).

**Figure 3.**
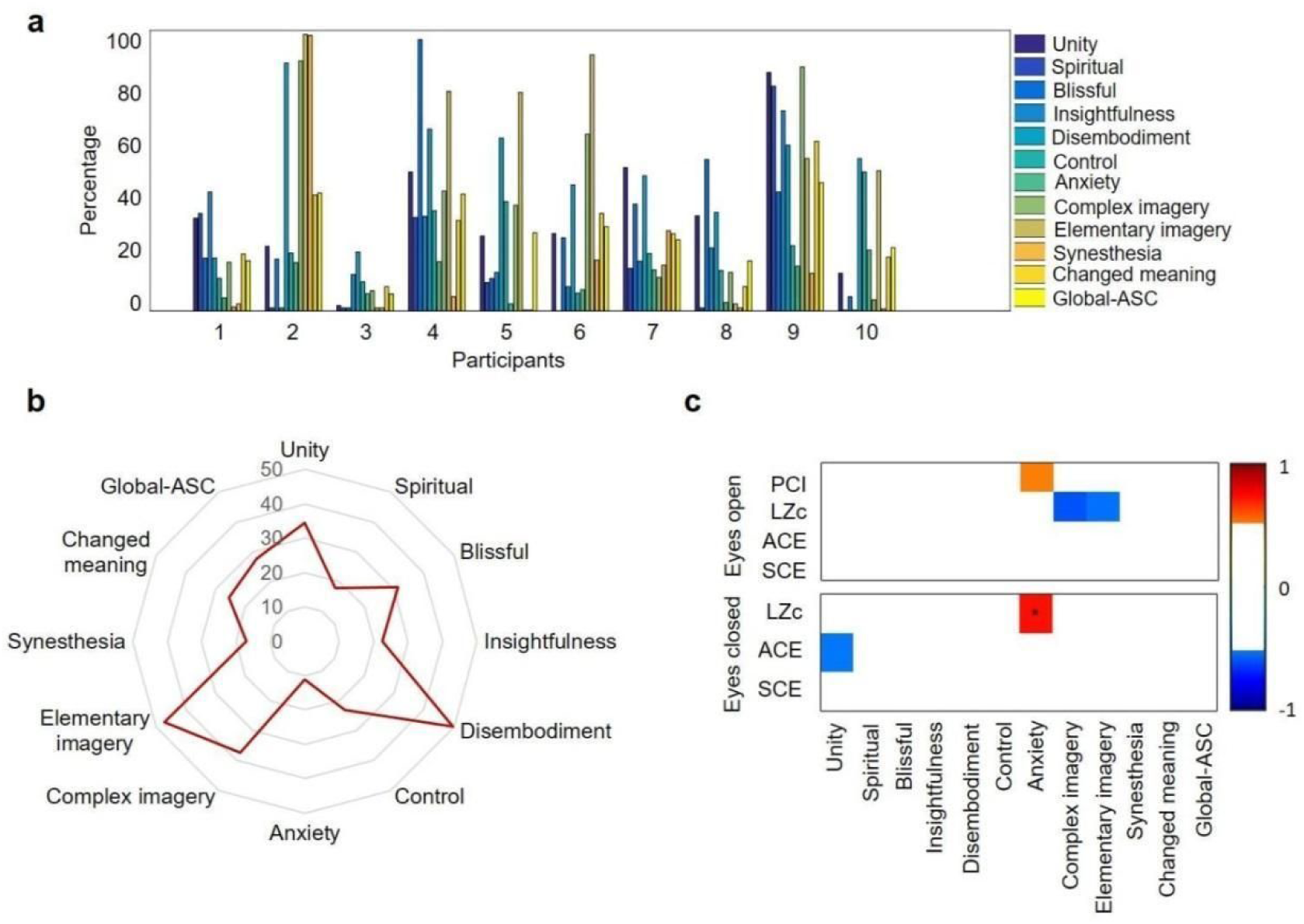
Phenomenology of sub-anaesthetic ketamine. **a)** Individual scores of the 11-Dimensional Altered States of Consciousness Rating Scale (11D-ASC) questionnaire, Global-ASC indicates the average score over all dimensions, **b**) Total mean scores for each dimension of the ASC questionnaire, **c**) correlation across signal diversity measures (difference scores between sub-anaesthetic ketamine and wakefulness) and ASC scores. Weak correlations (−0.5 < r < 0.5) are omitted (white) to only highlight strong correlations. Significance is indicated with a star (*).

## 3 Discussion

The main result of this study is the observation of significantly increased values of spontaneous EEG signal diversity measures in the ketamine-induced psychedelic state induced by sub-anaesthetic doses of ketamine, compared to the normal wakeful state. Moreover, we found a significant difference in spontaneous signal diversity in the eyes closed condition compared to eyes open during ketamine administration. In contrast, we observed no significant difference in PCI between the normal wakeful state and the ketamine-induced psychedelic state. As both PCI and spontaneous signal diversity have been seen to vary with the level of wakefulness in humans (Casali et al., 2013; Schartner, Carhart-Harris, et al., 2017; Schartner, Pigorini, et al., 2017; Schartner et al., 2015) these results suggest that PCI and spontaneous signal diversity measures may be sensitive to different aspects of conscious states.

### 3.1 Why were spontaneous signal diversity but not PCI values increased in the psychedelic state induced by sub-anaesthetic ketamine?

In both the normal wakeful state and sub-anaesthetic ketamine state, the spatiotemporal EEG responses to TMS perturbations seemed to contain fast, high-amplitude, and long-lasting waves of activity (**Figure 1**) resulting in no significant differences in PCI values (**Figure 2**). These results are consistent with the findings of Sarasso et al. (2015) where the PCI values measured during ketamine anaesthesia (i.e. with higher doses, causing a state of unresponsiveness after which all subjects reported vivid dreams) were similar to those measured during normal wakefulness, and high compared to PCI values obtained with other anaesthetics. In contrast, the spontaneous signal diversity measures, (LZ, ACE and SCE) were all significantly increased during sub-anaesthetic ketamine compared to the normal wakefulness condition before giving ketamine. This supports the findings from Schartner, Carhart-Harris, et al. (2017) that LZ and ACE, but not SCE, calculated from spontaneous MEG activity, increased significantly in the psychedelic states induced by ketamine, LSD, or psilocybin.

The difference between PCI and other signal diversity measures may seem unexpected, since previous studies have indicated that the measures of signal diversity (LZc, ACE, and SCE), based on spontaneous surface EEG and spontaneous intracranial EEG data, correspond quite well with PCI across conditions such as sleep (Schartner et al., 2015) and propofol anaesthesia (Schartner, Pigorini, et al., 2017). Since both our study and Schartner, Carhart-Harris, et al. (2017) found significant increases in the spontaneous signal diversity measures in the psychedelic state, one might have expected that also the PCI values should be increased similarly by sub-anaesthetic ketamine compared to normal wakefulness (Schartner, 2017). Why, then, did we find that sub-anaesthetic ketamine increased spontaneous signal diversity values but not PCI values?

One possible explanation for the observed differences could be that for PCI, complexity is computed from evoked EEG responses, while LZc, ACE and SCE are computed from spontaneous EEG. Both evoked and spontaneous signal diversity measures reflect differentiation because they are computed from diversity of patterns of activity over time and space (LZc and PCI), diversity over time over the most active channels (ACE), and diversity over time of synchronous channels (SCE) (Schartner et al., 2015). TMS evokes EEG responses that reflect causal interactions in the brain. In some states, the TMS-evoked activity may spread throughout the cortex, reflecting the degree of effective brain connectivity in that state (Massimini, Boly, Casali, Rosanova, & Tononi, 2009; Schartner, 2017). In contrast, signal diversity or complexity in spontaneous EEG is not necessarily strongly related to causal interactions, but could for example depend on degrees of independent vs. common driver inputs, thus being related to functional connectivity measures (Massimini et al., 2009; Schartner, 2017; Sitt et al., 2014). As such, PCI is considered to more closely reflect concurrent integration and differentiation by assessing the so-called “deterministic” responses to TMS, i.e. the responses that remain after averaging multiple trials (Casali et al., 2013). Crucially, PCI is designed to probe the general capacity of the brain to engage in complex causal interactions while being insensitive to the specific pattern of ongoing neural activity (Koch et al. 2016). This might explain the discrepancy between the PCI and spontaneous signal diversity results obtained here and by Schartner, Carhart-Harris, et al. (2017).

By design, spontaneous signal diversity measures are more sensitive than PCI to changes related to the complexity of the ongoing brain network activity. Our observations of the effects of eye closing seem to support this idea. We found that for all spontaneous signal diversity measures, there was a significant decrease when participants had their eyes closed compared to when they had open eyes. This may be related to the lack of visual input that increases synchronous alpha band activity (Barry, Clarke, Johnstone, Magee, & Rushby, 2007), thus ultimately affecting signal diversity. However, because drastically simplifying visual stimuli by closing the eyes might significantly reduce the complexity of the conscious content by reducing an aspect of the participant’s phenomenological experience, these findings suggest that spontaneous signal diversity may be more related to conscious content rather than conscious state. In contrast, having eyes closed does not seem to affect the TMS-EEG response (Rosanova et al., 2009, supplemental material) nor the PCI values during wakefulness (Casali et al., 2013, supplemental material). This suggests that PCI is less sensitive to changes in content (e.g. induced by removing visual aspects from phenomenal experience) within a global physiological state (here: being fully awake).

Moreover, spectral changes caused by sub-anaesthetic doses of ketamine may also be expected to influence signal diversity values. For example, sub-anaesthetic doses of ketamine have been shown to decrease alpha power in parallel with subjective ratings of dissociation of experience (Vlisides et al., 2018) as well as increase gamma power (de la Salle et al., 2016; Muthukumaraswamy et al., 2015). However, Schartner, Carhart-Harris, et al. (2017), using phase-shuffling normalization, found that the spectral profile changes seen in sub-anaesthetic ketamine could not explain the increased spontaneous signal diversity observed in the psychedelic condition.

Furthermore, the lack of significant change in PCI values in the psychedelic state compared to normal wakefulness (**Figures 1 and 2**) can be interpreted as low doses of ketamine causing so small changes in differentiation and integration that no change is detected in the TMS-evoked responses. Functional MRI studies have shown that hallucinogenic drugs produce an increased repertoire of activity patterns, thus increasing neural entropy compared to normal wakefulness (Lebedev et al., 2016; Tagliazucchi, Carhart-Harris, Leech, Nutt, & Chialvo, 2014) and reflecting increased differentiation of brain activity. This conclusion is supported by the changes in spontaneous EEG measures that we found here. However, it is possible that the increased signal diversity indicating neural differentiation may occur in parallel with a reduction of integration within the relevant brain networks, thus leading to a zero net change in PCI value. For example, Muthukumaraswamy et al. (2015) found that sub-anaesthetic doses of ketamine reduced frontal-to-parietal effective connectivity compared to normal wakefulness, and a similar decrease has been found in anaesthetic doses of ketamine (Lee et al., 2013). Yet, given the blocking effect of ketamine on NMDA-receptors and the widespread importance of these receptors throughout the cortex, it seems unlikely that reduced effective connectivity is limited to frontoparietal networks and it remains to be determined exactly how this decrease is related to overall cortical integration.

### 3.2 Relating signal diversity to phenomenological changes

The increase of signal diversity in the sub-anaesthetic ketamine condition may result from the brain state changes that cause the psychedelic experiences. Although all participants reported having had an effect of ketamine, the degree of subjective psychedelic effect, as measured by 11D-ASC, varied between participants (**Figure 3A**), which could be due to subjective differences in the quantification of the effect, different individual reactions to ketamine, or due to differences in administered ketamine dosage.

Similar to previous findings from placebo-controlled studies with sub-anaesthetic ketamine (Schmidt, Kometer, Bachmann, Seifritz, & Vollenweider, 2013; Studerus et al., 2010), the subscales *disembodiment* (feeling of dissociation between mind and body) and *elementary imagery* (changes in visual imagery with eyes closed) had the highest overall 11-ASC scores, while anxiety had the lowest overall scores (**Figure 3B**). Thus, the drug effects were largely as expected. The subscales *impaired cognition and control, anxiety* and *changed meaning of perception* had highest correlations with the signal diversity measures in the eyes open condition while *anxiety* had the highest correlation with eyes closed (**Figure 3C**, but note that anxiety was the dimension that showed the lowest overall score). In comparison, Schartner, Carhart-Harris, et al. (2017) found that increased spontaneous signal diversity in sub-anaesthetic ketamine was correlated with *overall intensity* of psychedelic experience as well as *ego-dissolution* and *vivid imagination*. These last two dimensions correspond to the subscales *disembodiment* and *complex imagery* (vivid complex visual patterns) from the 11D-ASC used in this study.

Furthermore, while the final version of this article was being prepared, D. Li and G.A. Mashour (2019) published a study on dose-dependent effects of ketamine. Using Hidden Markow modelling to classify spontaneous EEG signals into a discrete set of brain states, they found that the resulting Lempel-Ziv complexity was elevated during subanesthetic ketamine, while anesthetic doses caused alternating low and high complexity before stabilizing at a high level. Thus, using different methods, they arrived at results that seem to agree with ours for spontaneous EEG complexity during subanesthetic ketamine.

### 3.3 PCI and spontaneous signal diversity may reflect complementary aspects of consciousness

Behaviourally unresponsive states, such as the unresponsive wakefulness (“vegetative”) state, where subjects lack behaviourally verifiable intrinsic experiences, are associated with low PCI values compared to normal wakefulness (Casali et al., 2013; Casarotto et al., 2016). However, states in which subjects are behaviourally unresponsive but give delayed reports of having had vivid conscious experiences such as dreams during anaesthetic ketamine (Sarasso et al., 2015) and dreams during REM sleep (Casali et al., 2013), are associated with high PCI values similar to normal wakefulness. Therefore, PCI may reflect the brain’s capacity for sustaining experience per se, without differentiating whether the experience occurs with or without extrinsic awareness or ability to respond (Casali et al., 2013). Furthermore, since we did not find any difference in PCI comparing the sub-anaesthetic ketamine condition with normal wakefulness (all values in the ketamine condition were within the range of values typically reported for individuals in the awake state (Casarotto et al., 2016)), PCI may be more useful for differentiating between brain states with or without the capacity for sustaining consciousness, rather than measuring gradations of conscious content in wakeful experience.

Given that PCI values beyond the range found in normal wakefulness (i.e. up to 0.7) have so far not been measured (Casarotto et al., 2016), this poses the question of whether it is conceivable that there may exist brain states that give a measurably higher PCI value compared to the normal awake state. The psychedelic state, which has been described as involving “unconstrained cognition” (Carhart-Harris et al., 2012), has been considered a possible candidate. However, such changes in cognition may not be sufficient to exert a net change (increase) in PCI, suggesting some sort of “enhanced consciousness”. Rather, the psychedelic state might be a more “expansive” state in terms of “flexibility” in cognition compared to normal wakefulness, which could be reflected by increased signal diversity or entropy in the brain (Lebedev et al., 2016; Schartner, Carhart-Harris, et al., 2017; Tagliazucchi et al., 2014). Spontaneous signal diversity measures might therefore reflect increases in the complexity of conscious content. However, even though conscious content is modulated in the psychedelic state, this may not necessarily mean that the level of consciousness (i.e. level of arousal or wakefulness as in Laureys, Boly, Moonen, & Maquet, (2009) is altered. This also holds for having eyes open or closed, where PCI does not change, but signal diversity does. Furthermore, if LZc, ACE, and SCE only reflects the complexity of conscious content, and this complexity of content is separable from the capacity for consciousness, it makes sense that these measures correlate with PCI when comparing wakefulness with sleep (Schartner, Pigorini, et al., 2017) or anaesthesia (Schartner et al., 2015). This is because the conscious content normally appears to correlate with the level of wakefulness in these conditions (Laureys, Boly, Moonen, & Maquet, 2009).

### 3.4 Limitations

A limitation of our study is the moderate sample size. After 10 participants, we performed an interim analysis of the TMS-EEG data to evaluate continuation of the study and inclusion of more participants. As we did not find any significant differences in PCI values, and power tests could not predict significant differences nor correlations between PCI values and the ASC questionnaire with increased sample size, we decided not to continue. However, the sample size used in the current study is larger than in Sarasso et al. (2015) where no difference in PCI was found for anaesthetic doses of ketamine compared to normal wakefulness, and our spontaneous signal diversity results are similar to Schartner, Carhart-Harris, et al. (2017) where number of participants was higher.

In addition, we administered ketamine to the participants as a continuous infusion, gradually increasing the dose until the participants reported an effect of the drug instead of giving a pre-defined bolus and maintenance dose. The goal was to ensure that all participants had comparable drug effects on subjective experiences. However, as the infusion was not always increased in equal steps (**Figure S3**) there were different continuous infusion rates and total doses of ketamine for each participant (particularly for the first 3 participants, where the doses were increased in smaller steps and over longer time than for the other 7 participants. A bolus dose of ketamine (de la Salle et al., 2016; Driesen et al., 2013) could have avoided this complication, and allowed for placebo control, but would not have ensured comparable subjective drug effects for all participants. Although the differences in infusion rate and dose may have affected PCI and 11D-ASC scores, we found no significant correlation between total and continuous dosage (infusion rate or total dose), nor between total dose and subjective experience. Although, we cannot exclude the possibility that expectation bias affected the participants’ evaluation, our study confirmed findings from similar studies that included placebo control (Schartner, Carhart-Harris, et al., 2017). Placebo control is anyway difficult for psychedelic states when it is particularly easy for a participant to determine whether they got an active or inactive compound.

### 3.5 Future directions

Further exploring how different proposed markers of consciousness are affected in conditions where phenomenal content is altered may help us understand which aspects of consciousness the markers are most closely linked to. Understanding how these markers respond to conditions that induce changes in phenomenology, by pharmacology (e.g. anaesthetics or psychedelics), pathology (e.g. in patients with psychiatric disorders or disorders of consciousness), or physiological brain states (e.g. sleep, in “flow states”, or sensory deprivation) can help us understand the relations between functional, dynamical, or structural properties of the brain and conscious experience. Thus, further testing, comparing, and contrasting promising neurophysiological markers of consciousness in conditions where phenomenology is altered and mapping out the relation between observed changes in the markers and reported changes in the structure of experience, may help identifying sets of complementary markers that are sensitive to distinct aspects of conscious experience. Furthermore, it may help deciding whether unnecessarily complex measures can be exchanged for a set of simpler markers for some practical purposes.

## 4 Conclusion

In this study we investigated whether TMS-evoked (PCI) and spontaneous EEG signal diversity measures (LZc, ACE, and SCE) are affected by sub-anaesthetic doses of ketamine causing a psychedelic state, compared to the normal wakeful state. We found no significant change in PCI when comparing sub-anaesthetic ketamine and normal wakefulness. However, we did find significant increases in spontaneous EEG signal diversity when participants were under the influence of sub-anaesthetic doses of ketamine compared to normal wakefulness. We also found that the spontaneous signal diversity measures were significantly lower with eyes closed than with eyes open, both during normal wakefulness and under the effect of sub-anaesthetic ketamine doses. Furthermore, we found correlations between changes in spontaneous signal diversity measures and subjective ratings of phenomenological experience. These results suggest that spontaneous and evoked measures of EEG signal diversity may reflect distinct, complementary aspects of changes in brain function related to altered states of consciousness. Spontaneous EEG measures may thus capture properties related to the content of consciousness, while evoked measures (PCI) may index the system’s capacity for consciousness.

## 5 Acknowledgements

We are thankful to Marcello Massimini, Silvia Casarotto, and Matteo Fecchio for providing computer software and insightful, technical help, and support regarding the PCI measurements, and to M. Massimini also for comments on the manuscript. We also thank Pål G. Larsson for technical advice regarding EEG, Franz Vollenveider for advice regarding ketamine administration, Morten Engstrøm for valuable feedback during the writing of the manuscript, Brita Noorland for help with ketamine administration, and Anikó Kuztor for helping with the data acquisition and analysis. This study was supported by the European Union’s Horizon 2020 research and innovation programme under grant agreement 7202070 (Human Brain Project (HBP)) and the Norwegian Research Council (NRC: 262950/F20 and 214079/F20).

## 7 Supplementary material

**Figure S1.**
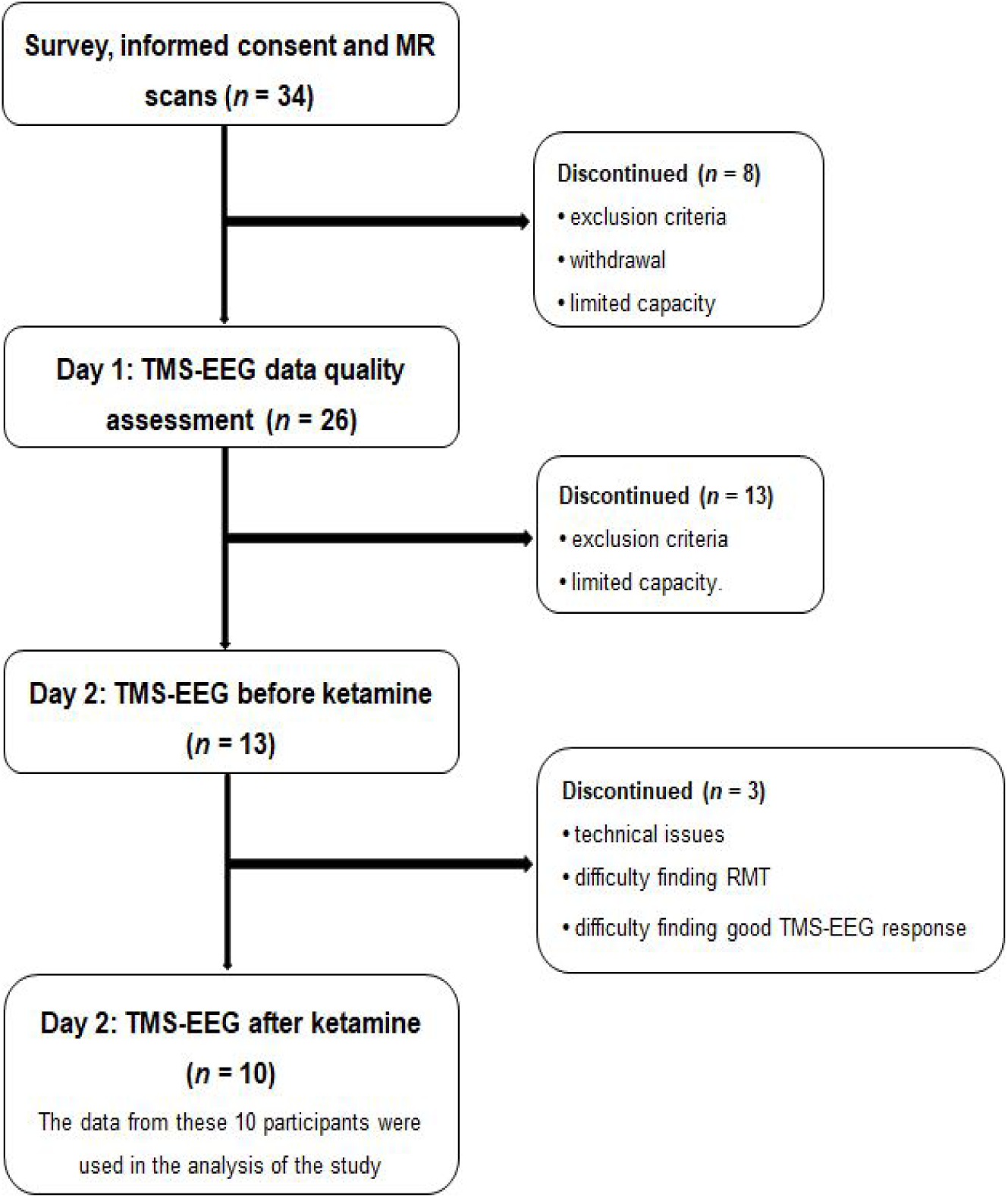
Flowchart of participation in the different stages of the study. The number of participants included (left) or discontinued (right) at each stage of the study is indicated with *n*. In the end 10 participants completed the study.

**Figure S2.**
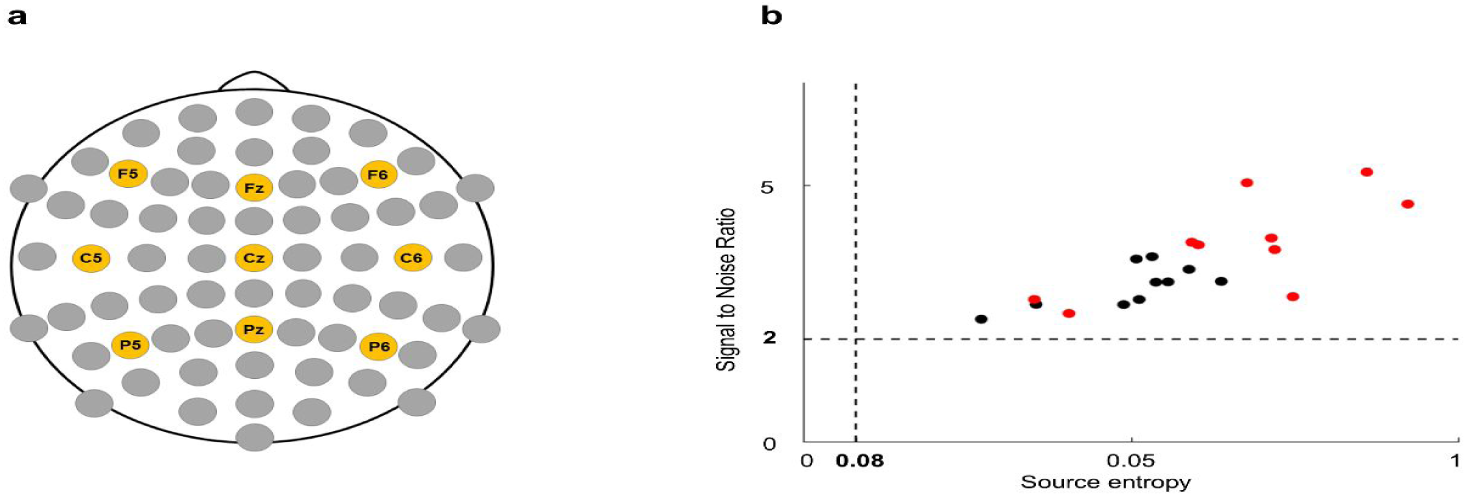
Electrode positions, signal-to-noise ratio (SNR), and source entropy (H). **a**) Spontaneous signal diversity was calculated from 9 channels (orange) distributed across the scalp: F5, Fz, F6, C6, Cz, C6, P5, Pz, and P6. **b**) Signal-to-noise ratio (SNR) and source entropy (H) for all TMS-EEG measurements and participants. The black dots are participants during normal wakefulness and the red dots are participants during sub-anaesthetic ketamine. The source entropy of the data was above 0.08 and all data showed a high SNR (> 2). None of the data were excluded based on too low source entropy.

**Figure S3.**
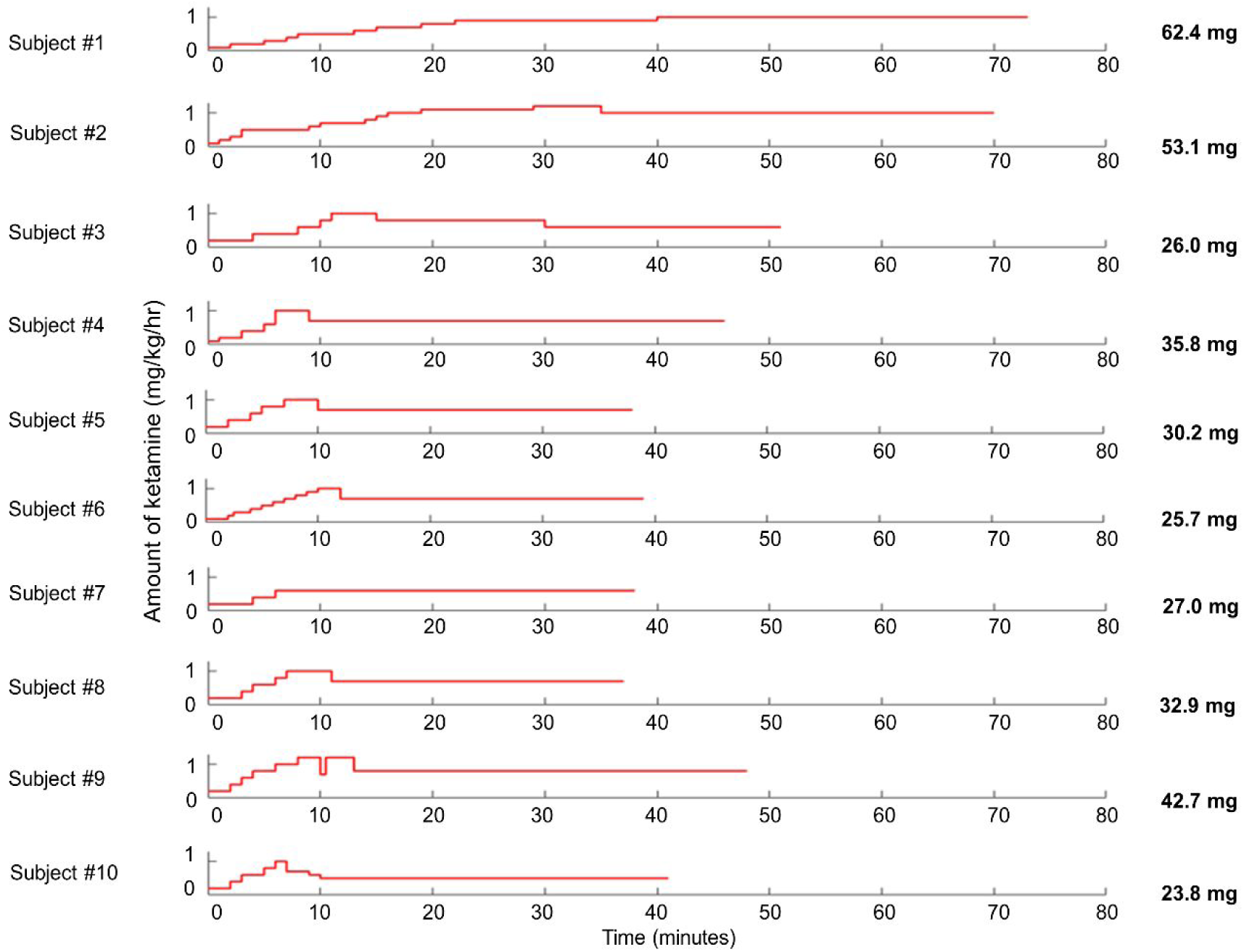
Ketamine administration. The plots show the time course of ketamine infusion rates, between 0.1 and 1.2 mg/kg/hr for the ten subjects. The infusion was eventually stabilized (0.1 – 1.0 mg/kg/hr) at a constant rate in the final period, during which, all the TMS-EEG data used in the study were collected. The total amount of ketamine given is shown to the right.

**Figure S4.**
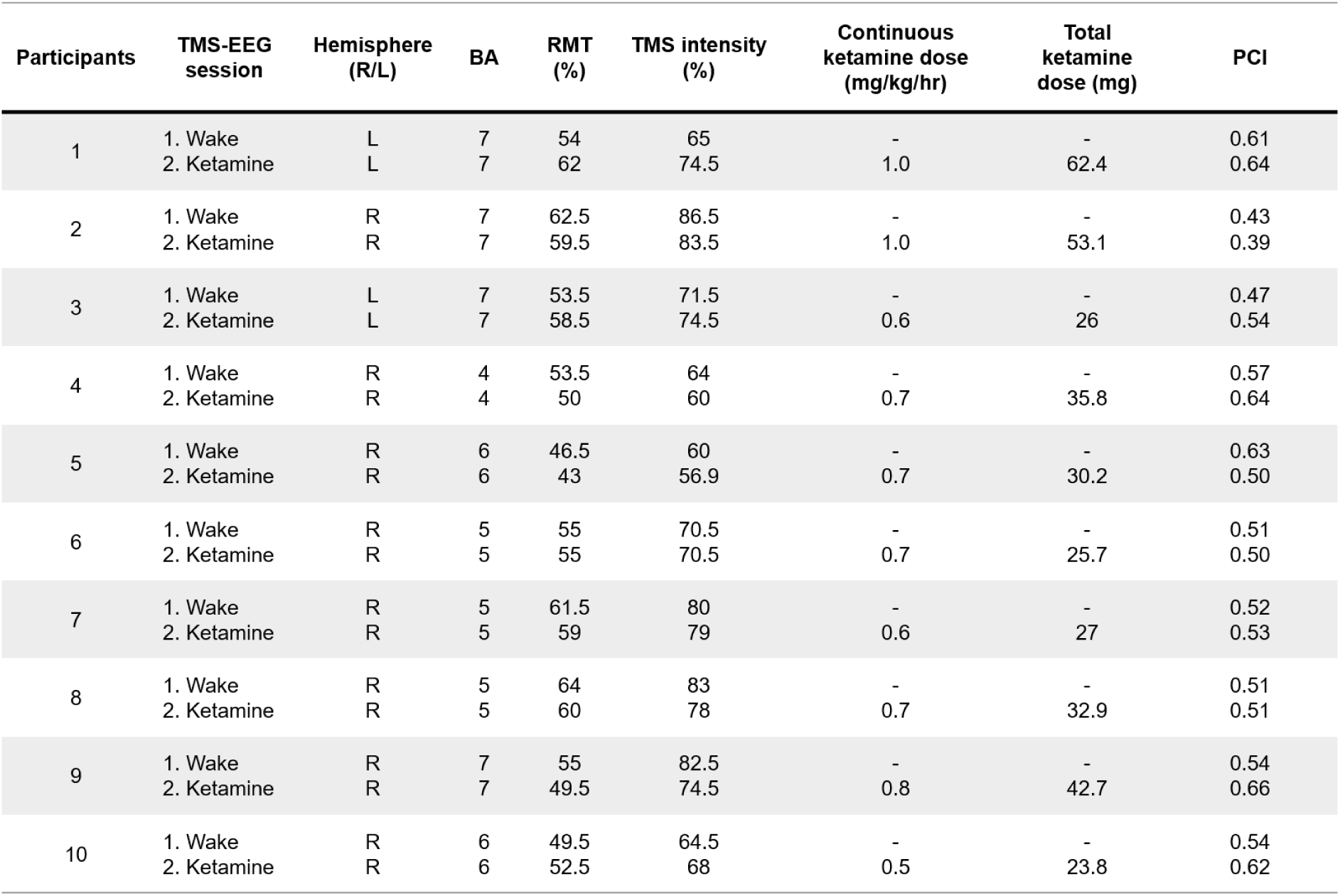
**Stimulation parameters, ketamine doses and PCI values** for the TMS-EEG sessions. PCI values in bold were used in the present study. **R** = right, **L** = left, **BA** = Brodmann Area, **%** = percentage of maximal stimulator output.

## References

Barry, R. J., Clarke, A. R., Johnstone, S. J., Magee, C. A., & Rushby, J. A. (2007). EEG differences between eyes-closed and eyes-open resting conditions. Clin Neurophysiol, 118(12), 2765–2773. doi:10.1016/j.clinph.2007.07.028

Bayne, T., & Carter, O. (2018). Dimensions of consciousness and the psychedelic state. Neuroscience of Consciousness, 2018(1), iy008–niy008. doi:10.1093/nc/niy008

Bayne, T., Hohwy, J., & Owen, A. M. (2016). Are There Levels of Consciousness? Trends in cognitive sciences, 20(6), 405–413. doi:10.1016/j.tics.2016.03.009

Boly, M., Seth, A., Wilke, M., Ingmundson, P., Baars, B., Laureys, S., … Tsuchiya, N. (2013). Consciousness in humans and non-human animals: recent advances and future directions. Frontiers in Psychology, 4(625). doi:10.3389/fpsyg.2013.00625

Bowdle, T. A., Radant, A. D., Cowley, D. S., Kharasch, E. D., Strassman, R. J., & Roy-Byrne, P. P. (1998). Psychedelic effects of ketamine in healthy volunteers: relationship to steady-state plasma concentrations. Anesthesiology, 88(1), 82–88.

Carhart-Harris, R. L., Erritzoe, D., Williams, T., Stone, J. M., Reed, L. J., Colasanti, A., … Nutt, D. J. (2012). Neural correlates of the psychedelic state as determined by fMRI studies with psilocybin. Proceedings of the National Academy of Sciences, 109(6), 2138–2143. doi:10.1073/pnas.1119598109

Casali, A. G., Gosseries, O., Rosanova, M., Boly, M., Sarasso, S., Casali, K. R., … Massimini, M. (2013). A Theoretically Based Index of Consciousness Independent of Sensory Processing and Behavior. Science Translational Medicine, 5(198), 198ra105.

Casarotto, S., Comanducci, A., Rosanova, M., Sarasso, S., Fecchio, M., Napolitani, M., … Massimini, M. (2016). Stratification of unresponsive patients by an independently validated index of brain complexity. Annals of Neurology, 80(5), 718–729. doi:10.1002/ana.24779

de la Salle, S., Choueiry, J., Shah, D., Bowers, H., McIntosh, J., Ilivitsky, V., & Knott, V. (2016). Effects of Ketamine on Resting-State EEG Activity and Their Relationship to Perceptual/Dissociative Symptoms in Healthy Humans. Frontiers in Pharmacology, 7, 348. doi:10.3389/fphar.2016.00348

Di Lazzaro, V., Oliviero, A., Profice, P., Pennisi, M. A., Pilato, F., Zito, G., … Tonali, P. A. (2003). Ketamine increases human motor cortex excitability to transcranial magnetic stimulation. J Physiol, 547(Pt 2), 485–496. doi:10.1113/jphysiol.2002.030486

Di, X., Gohel, S., Kim, E., & Biswal, B. (2013). Task vs. rest—different network configurations between the coactivation and the resting-state brain networks. Frontiers in Human Neuroscience, 7(493). doi:10.3389/fnhum.2013.00493

Dittrich, A. (1998). The standardized psychometric assessment of altered states of consciousness (ASCs) in humans. Pharmacopsychiatry, 31 Suppl 2, 80–84. doi:10.1055/s-2007-979351

Domino, E. F., Chodoff, P., & Corssen, G. (1965). Pharmacologic effects of CI-581, a new dissociative anesthetic, in main. Clin Pharmacol Ther, 6, 279–291.

Driesen, N. R., McCarthy, G., Bhagwagar, Z., Bloch, M., Calhoun, V., D’Souza, D. C., … Krystal, J. H. (2013). Relationship of resting brain hyperconnectivity and schizophrenia-like symptoms produced by the NMDA receptor antagonist ketamine in humans. Mol Psychiatry, 18(11), 1199–1204. doi:10.1038/mp.2012.194

Farnes, N., Juel, B.E., Nilsen, A.S., Romundstad, L.G., Larsson, P.G., Engstrøm, M., Storm, J.F. (2017a). An index of consciousness, Perturbational Complexity Index (PCI), compared between normal awake state and psychedelic state induced by sub-anaesthetic doses of ketamine. Poster at Nordic Neuroscience 2017. Abstract available from http://nordicneuroscience.org/2017/wp-content/uploads/2016/06/Abstract-Book-Nordic-Neuroscience-2017.pdf

Farnes, N., Juel, B.E., Nilsen, A.S., Romundstad, L.S., Larsson, P.G., Engstrøm, M. & Storm, J.F., (2017b) Consciousness and Cortical Connectivity in Subanaesthetic Ketamine Measured by TMS and EEG. In Simidjievski, N., Santuy, A., Menzel, M., Tanevski, J., Karasenko, V., Mahfoud, T. … Saria, A. 1st HBP Student Conference - Transdisciplinary Research Linking Neuroscience, Brain Medicine and Computer Science. Poster presented at Human Brain Project Student Conference, Vienna, 8-10 february (pp. 92-94). Frontiers Media SA. Abstract available from https://www.frontiersin.org/books/1st_HBP_Student_Conference_-_Transdisciplinary_Research_Linking_Neuroscience_Brain_Medicine_and_Com/1532

Farnes, N., Juel, B.E., Nilsen, A.S., Romundstad, Storm, J.F. (2019). Increased signal diversity/complexity of spontaneous EEG in humans given sub-anaesthetic doses of ketamine. BiorXiv. https://doi.org/10.1101/508697

Ferrarelli, F., Riedner, B. A., Peterson, M. J., & Tononi, G. (2015). Altered prefrontal activity and connectivity predict different cognitive deficits in schizophrenia. Hum Brain Mapp, 36(11), 4539–4552.

Gallimore, A. R. (2015). Restructuring consciousness –the psychedelic state in light of integrated information theory. Frontiers in Human Neuroscience, 9(346). doi:10.3389/fnhum.2015.00346

Gress, D. R. (2009). Plum and Posner’s diagnosis of stupor and coma, 4th edition. Neurology, 72(3), 295–295. doi:10.1212/01.wnl.0000339492.66776.fc

Hamidi, M., Slagter, H. A., Tononi, G., & Postle, B. R. (2010). Brain responses evoked by high-frequency repetitive TMS: An ERP study. Brain Stimul, 3(1), 2–17. doi:10.1016/j.brs.2009.04.001

Hjorth, B. (1975). An on-line transformation of EEG scalp potentials into orthogonal source derivations. Electroencephalography and Clinical Neurophysiology, 39(5), 526–530. doi:https://doi.org/10.1016/0013-4694(75)90056-5

Ilmoniemi, R. J., & Kicic, D. (2010). Methodology for combined TMS and EEG. Brain Topogr, 22(4), 233–248. doi:10.1007/s10548-009-0123-4

Kaspar, F., & Schuster, H. G. (1987). Easily calculable measure for the complexity of spatiotemporal patterns. Physical Review A, 36(2), 842–848. doi:10.1103/PhysRevA.36.842

Koch, C., Massimini, M., Boly, M., & Tononi, G. (2016). Neural correlates of consciousness: progress and problems. Nat Rev Neurosci, 17(5), 307–321. doi:10.1038/nrn.2016.22

Laureys, S., Boly, M., Moonen, G., & Maquet, P. (2009). Two dimensions of consciousness: arousal and awareness. Encycl Neurosci, 2, 1133–1142.

Lebedev, A. V., Kaelen, M., Lovden, M., Nilsson, J., Feilding, A., Nutt, D. J., & Carhart-Harris, R. L. (2016). LSD-induced entropic brain activity predicts subsequent personality change. Hum Brain Mapp, 37(9), 3203–3213. doi:10.1002/hbm.23234

Lee, U., Ku, S., Noh, G., Baek, S., Choi, B., & Mashour, G. A. (2013). Disruption of Frontal-Parietal Communication by Ketamine, Propofol, and Sevoflurane. Anesthesiology, 118(6), 1264–1275. doi:10.1097/ALN.0b013e31829103f5

Lempel, A., & Ziv, J. (1976). On the complexity of finite sequences. IEEE Transactions on information theory, 22(1), 75–81.

Li, D., & Mashour, G. A. (2019). Cortical dynamics during psychedelic and anesthetized states induced by ketamine. NeuroImage, 196, 32–40.

Massimini, M., Boly, M., Casali, A., Rosanova, M., & Tononi, G. (2009). A perturbational approach for evaluating the brain’s capacity for consciousness. In N. D. S. Steven Laureys & M. O. Adrian (Eds.), Progress in Brain Research (Vol. Volume 177, pp. 201–214): Elsevier.

Muthukumaraswamy, S. D., Shaw, A. D., Jackson, L. E., Hall, J., Moran, R., & Saxena, N. (2015). Evidence that Subanesthetic Doses of Ketamine Cause Sustained Disruptions of NMDA and AMPA-Mediated Frontoparietal Connectivity in Humans. J Neurosci, 35(33), 11694–11706. doi:10.1523/jneurosci.0903-15.2015

Nagel, T. (1974). What Is It Like to Be a Bat? The Philosophical Review, 83(4), 435–450. doi:10.2307/2183914

Oizumi, M., Albantakis, L., & Tononi, G. (2014). From the Phenomenology to the Mechanisms of Consciousness: Integrated Information Theory 3.0. PLOS Computational Biology, 10(5), e1003588. doi:10.1371/journal.pcbi.1003588

Rogasch, N. C., Sullivan, C., Thomson, R. H., Rose, N. S., Bailey, N. W., Fitzgerald, P. B., … Hernandez-Pavon, J. C. (2017). Analysing concurrent transcranial magnetic stimulation and electroencephalographic data: A review and introduction to the open-source TESA software. Neuroimage, 147, 934–951. doi:http://dx.doi.org/10.1016/j.neuroimage.2016.10.031

Rosanova, M., Casali, A., Bellina, V., Resta, F., Mariotti, M., & Massimini, M. (2009). Natural Frequencies of Human Corticothalamic Circuits. The Journal of Neuroscience, 29(24), 7679–7685. doi:10.1523/jneurosci.0445-09.2009

Rossi, S., Hallett, M., Rossini, P. M., Pascual-Leone, A., & Safety of T. M. S. Consensus Group, T. (2009). Safety, ethical considerations, and application guidelines for the use of transcranial magnetic stimulation in clinical practice and research. Clinical Neurophysiology, 120(12), 2008–2039. doi:10.1016/j.clinph.2009.08.016

Rossini, P. M., Burke, D., Chen, R., Cohen, L. G., Daskalakis, Z., Di Iorio, R., … Ziemann, U. (2015). Non-invasive electrical and magnetic stimulation of the brain, spinal cord, roots and peripheral nerves: Basic principles and procedures for routine clinical and research application. An updated report from an I.F.C.N. Committee. Clinical Neurophysiology, 126(6), 1071–1107. doi:http://dx.doi.org/10.1016/j.clinph.2015.02.001

Rotenberg, A., Horvath, J. C., & Pascual-Leone, A. (2014). Transcranial Magnetic Stimulation (1 ed. Vol.89): Humana Press.

Sarasso, S., Boly, M., Napolitani, M., Gosseries, O., Charland-Verville, V., Casarotto, S., … Massimini, M. (2015). Consciousness and Complexity during Unresponsiveness Induced by Propofol, Xenon, and Ketamine. Current Biology, 25(23), 3099–3105. doi:10.1016/j.cub.2015.10.014

Schartner, M. M. (2017). On the relation between complex brain activity and consciousness. (Doctoral), University of Sussex. Retrieved from http://sro.sussex.ac.uk/67112/

Schartner, M. M., Carhart-Harris, R. L., Barrett, A. B., Seth, A. K., & Muthukumaraswamy, S. D. (2017). Increased spontaneous MEG signal diversity for psychoactive doses of ketamine, LSD and psilocybin. Scientific Reports, 7, 46421. doi:10.1038/srep46421

Schartner, M. M., Pigorini, A., Gibbs, S. A., Arnulfo, G., Sarasso, S., Barnett, L., … Barrett, A. B. (2017). Global and local complexity of intracranial EEG decreases during NREM sleep. Neuroscience of Consciousness, 3(1), iw022–niw022. doi:10.1093/nc/niw022

Schartner, M. M., Seth, A., Noirhomme, Q., Boly, M., Bruno, M.-A., Laureys, S., & Barrett, A. (2015). Complexity of Multi-Dimensional Spontaneous EEG Decreases during Propofol Induced General Anaesthesia. PLOS ONE, 10(8), e0133532. doi:10.1371/journal.pone.0133532

Schmidt, A., Kometer, M., Bachmann, R., Seifritz, E., & Vollenweider, F. (2013). The NMDA antagonist ketamine and the 5-HT agonist psilocybin produce dissociable effects on structural encoding of emotional face expressions. Psychopharmacology, 225(1), 227–239. doi:10.1007/s00213-012-2811-0

Sitt, J. D., King, J.-R., El Karoui, I., Rohaut, B., Faugeras, F., Gramfort, A., … Naccache, L. (2014). Large scale screening of neural signatures of consciousness in patients in a vegetative or minimally conscious state. Brain, 137(8), 2258–2270. doi:10.1093/brain/awu141

Studerus, E., Gamma, A., & Vollenweider, F. X. (2010). Psychometric Evaluation of the Altered States of Consciousness Rating Scale (OAV). PLOS ONE, 5(8), e12412. doi:10.1371/journal.pone.0012412

Tagliazucchi, E., Carhart-Harris, R., Leech, R., Nutt, D., & Chialvo, D. R. (2014). Enhanced repertoire of brain dynamical states during the psychedelic experience. Hum Brain Mapp, 35(11), 5442–5456. doi:10.1002/hbm.22562

Tononi, G., Boly, M., Massimini, M., & Koch, C. (2016). Integrated information theory: from consciousness to its physical substrate. Nature Reviews Neuroscience, 17, 450. doi:10.1038/nrn.2016.44

Tononi, G., & Edelman, G. M. (1998). Consciousness and Complexity. Science, 282(5395), 1846.

Vlisides, P. E., Bel-Bahar, T., Lee, U., Li, D., Kim, H., Janke, E., … & Picton, P. (2017). Neurophysiologic Correlates of Ketamine Sedation and Anesthesia A High-density Electroencephalography Study in Healthy Volunteers. Anesthesiology: The Journal of the American Society of Anesthesiologists, 127(1), 58–69.

